# Flexibility in the face of climate change? A rapid and dramatic shift toward later spring migration in Hudsonian Godwits (*Limosa haemastica*)

**DOI:** 10.1101/2024.04.30.591937

**Authors:** Lauren Puleo, Feipeng Huang, Maria Stager, Nathan R. Senner

## Abstract

With rapid environmental change, shifts in migration timing are vitally important for maintaining population stability and have been widely documented. However, little remains known about *how* migrants are driving these shifts and what factors may influence the effective utilization of these strategies, limiting our ability to accurately assess species- and population-level vulnerability to climate change. The Hudsonian godwit (*Limosa haemastica*) is an extreme long-distance migratory shorebird that has (1) previously shifted its population-level migration timing and (2) exhibits sex-specific morphological differences. Therefore, we combined over a decade of light-level geolocator tracking data from a single breeding population with a historical predictive model to assess on-going shifts in migration timing while determining the time-shifting strategies utilized by each sex. Surprisingly, we found that godwit departure and arrival timing rapidly shifted 6 days later from 2010-2023 with no differences in timing between the sexes. Despite this change in migration timing, the population has maintained an average migratory duration of 24 days, suggesting that godwits are driving shifts in arrival timing entirely by shifting their nonbreeding ground departure, something rarely documented in long-distance migrants. Yet, we also found that godwits are not shifting their migration timing in the direction predicted by our model, providing evidence that this response may not be adaptive. These results emphasize the urgent need for a more holistic approach to assessing the relative vulnerability of migratory species and the adaptiveness of changes in migration timing.

## INTRODUCTION

As climate change alters the phenology of the resources on which migratory species and their offspring rely^1^, the ability to adjust migratory timing is vital for maintaining individual fitness and stable populations ^2^. As a result, large-scale changes in migratory timing, particularly advancements during spring, have been widely documented and the degree to which populations have adjusted their timing remains a critical measure when assessing current and future vulnerabilities to climate change ^3,4^. The rate at which migratory timing is shifting, however, remains highly variable at the population- and species-levels ^5^. This suggests the need to not only document these modifications, but also investigate the factors that influence an individual’s or population’s ability to utilize different time-shifting strategies – an often-overlooked component of large-scale migration phenology studies ^6^.

Long-distance migrants, in particular, face an array of climatic and non-climatic factors that are likely to influence their ability to adequately respond to phenological changes. For instance, long-distance migrants are thought to inherently rely on invariant photoperiodic cues to initiate nonbreeding ground departure, which would make this an unlikely point in the annual cycle to catalyze a shift in migratory timing ^7,8^. Recent studies have shown, though, that some long-distance migrants can use information gathered during previous migrations to adjust their departure timing ^9^, but it remains unclear whether these shifts can lead to advancements in breeding ground arrival ^10^. In general, the success of this strategy depends on the interannual variation in breeding ground conditions experienced by a population and the anomalousness of conditions within a given year ^11^.

Utilizing *en route* decisions may be more viable for long-distance migrants, as they may be able to encounter increasingly reliable cues about breeding ground conditions along their migration route ^12^. Yet, decisions made *en route*, such as spending less time at stopover sites or skipping stopover sites altogether, may reduce time needed for refueling – potentially affecting males and females differentially because of divergent, sex-specific goals during migration. For instance, males are likely under strong time selection to secure breeding territories ^13^ and avoid the fitness consequences of divorce ^14,15^, whereas females have the added pressure of accruing the resources needed for reproduction ^16^. Thus, despite leading to earlier breeding ground arrival, a reduction in refueling time can result in reduced female energetic condition upon arrival, delaying reproduction and failing to alleviate climate-induced phenological mismatches ^17^. The potential for females to face such tradeoffs between arrival timing and condition emphasizes that sex-specific goals can affect the utilization and effectiveness of time-shifting strategies for each sex, as well as constrain an entire population’s response to phenological shifts ^18^. Ultimately, an increased understanding of sex-specific differences in timing and the time-shifting strategies utilized could further elucidate the potential for climate change induced reversible-state effects to affect reproduction ^19^ a critical gap in our understanding that is inhibiting comprehensive vulnerability assessments of migratory species.

Hudsonian godwits (*Limosa haemastica*; hereafter ‘godwits’) are extreme long-distance migrants that spend the nonbreeding season in southern South America and breed in sub-arctic North America ^20^. Despite their predicted vulnerability to climatic change, the godwit population breeding in Beluga, Alaska appears to be using knowledge from previous migrations to effectively track long-term slow, linear, climate-driven shifts in phenology ^11^. As a result, Beluga-breeding godwits advanced both their spring migratory and reproductive timing by 9 days between 1974-2010 ^11,21^. Nonetheless, in anomalously early and warm years, godwits experience phenological mismatches that significantly reduce reproductive success ^22^. Godwits also exhibit high levels of mate fidelity and in some cases, rapid clutch initiation following migration ^20^, suggesting that sex-specific goals may affect the strategies each sex can use to adjust their migratory timing. Yet, it remains unknown whether godwits have been continuing to effectively track more recent, increasingly variable, climatic trends, and *how* individuals might be achieving these shifts.

Here we used long-term geolocation tracking data to assess on-going changes in godwit migratory timing by investigating **(1)** nonbreeding ground departure timing **(2)** breeding ground arrival timing and **(3)** migration duration, while investigating potential sex-specific differences at each stage. With parameters from a historical model ^11^, we then predicted godwit breeding ground first arrival dates using eBird and environmental data to compare with first-arrival dates from our tracking data and determine if godwits have continued to track recent shifts in the climate regime. We expected that godwit breeding ground arrival timing would continue to trend earlier as we had found previously. Additionally, because we expected males to be under strong time selection, we predicted that males would utilize *en route* time-minimizing strategies, reducing migration duration over time and driving population-level trends in breeding ground arrival timing. However, while we still expected to see females shift their spring migratory timing earlier to some degree, we predicted that there would be little reduction, if any, in migration duration because of the added pressure to accrue the resources needed for egg production *en route*. Our study illuminates the time-shifting strategies used by individuals to alter migratory timing and determines if demographic factors, such as sex, can influence the utilization and effectiveness of such strategies. Ultimately, our work will add to the growing body of knowledge regarding the ability of migratory species to flexibly respond to a changing climate.

## METHODS

### (1) Godwit capture and tracking

We monitored godwits in Beluga (61°21′N, 151°03′W) during the breeding season (May – July) from 2009-2023 and tracked the migrations of these breeding birds using light-level geolocators during two separate periods: 2009-2012 (British Antarctic Survey Mk14 geolocators) ^23^ and 2020-2023 (Migrate Technology Ltd. Intigeo geolocators). Tracked individuals were captured on the nest using mist-nets during incubation and their sex determined by plumage and morphological measurements ^20^. Each individual was fit with a unique alpha-numeric USGS metal band on the right tibia, in addition to a plastic color cohort band and individually specific green leg flag with a geolocator attached to it on the left tibia. In subsequent breeding seasons, geolocators were deployed on previously tagged individuals that were re-captured and had their old geolocator replaced, as well as previously unmarked individuals. In total, we retrieved 81 geolocators, 52 (female=21 & male=31) from 2009-2012 and 29 (female=15 & male=14) from 2020-2023.

### (2) Data filtering

Raw light data for each individual was extracted using BASTrack (2009-2012) or IntigeoIF (2020-2023), and then analyzed in the R Programing Environment (v.4.2.2, R Core Development Team 2023) using the packages ‘GeoLight’ (v.2.0.1) ^24^ and ‘FLightR’ (v.0.5.3) ^25^ following Lisovski et al. ^26^. Twilights (i.e., sunrises and sunsets) were annotated and outliers that were incorrectly classified (e.g., periods where the geolocator was shaded because of incubation) were identified by visual inspection of the raw light data and either adjusted manually or deleted. Filtered twilights were then calibrated in ‘FLightR’ using the known deployment location coordinates, with individually distinct calibration periods chosen based on when an individual’s geolocator was deployed or through an assessment of the raw light data. We extracted migration schedules using a maximum flight distance between two consecutive twilight periods of 1900 km and a probability cutoff of 0.2. Mean latitudes and longitudes were used to designate stopover locations. The estimated nonbreeding ground departure and breeding ground arrival dates were selected from the median quartile and transformed into Julian dates for analysis. In some cases, geolocators failed to capture data during the migration window, raw light data exhibited timing drift, or location estimations produced in ‘FLightR’ were poor (e.g., location estimates falling outside of the species’ known range) despite using the above parameters. This resulted in an exclusion of 16 geolocators from subsequent analysis. Of the 65 remaining geolocators, five had data from two migrations, giving us a total of 70 tracks from round-trip migrations, with 19 individuals providing tracks for more than one migration (hereafter, ‘multi-year individuals’).

Accurately identifying arrival and departure dates with geolocators is notoriously difficult ^25^. We therefore also used the wet/dry data collected by the geolocators to corroborate our estimates of godwit departure and arrival timing throughout their migrations. British Antarctic Survey geolocators (2009-

2012) recorded conductivity measurements every 10 minutes, with a scale ranging from 0-200, where measurements of 0 represent complete dryness. Migrate Technology Ltd. geolocators (2020-2023) recorded how many times the logger was wet every 30 seconds over each 4-hour period (measures of 0-480), as well as the maximum conductivity value recorded during that 4-hour period (scale of 0-127). For species that take multi-day non-stop migratory flights like godwits, distinct periods of dryness (i.e., conductivity or wetness measures of 0) can be used to accurately gauge when an individual departed from one location and arrived at another ^27^. The accuracy of this method is further strengthened by godwit foraging behavior, as they feed primarily on coastlines or in wetlands, ensuring submersion of the geolocator when an individual has ceased flying.

When timing estimates varied between ‘FLightR’ and our wet/dry data, we utilized the first day of a recorded dry block as the ‘true’ departure date and the first day of a recorded wet block following a dry block as the ‘true’ arrival date. Nonetheless, timing estimates never varied by more than 3 days between our methods. For 5 tracks used in the final analysis, wet/dry data were not obtained – in most cases this was a result of tags that were retrieved with 2 years of movement data and for which wet/dry data was only captured during the first year the geolocator was deployed. In these instances, ‘FLightR’ location and timing estimates were used without wet/dry corroboration. To account for the potential reduction in accuracy in departure timing estimation for these individuals, we chose the latest possible departure date identified by ‘FlightR’ as the ‘true’ departure and the earliest possible arrival date as the ‘true’ arrival to the breeding grounds.

### (3) Analysis

All models were run in the R Programming Environment using the package ‘*lme4’* ^28^and their statistical significance evaluated using the package ‘*lmertest’* ^29^.

#### (a) Population- and individual-level analysis

First, we investigated shifts in breeding ground arrival and nonbreeding ground departure within the two distinct tracking periods (2009-2012, tracks=40; 2020-2023, tracks=30). To do this, we used linear mixed-effects models with year as the predictor variable and individual as a random effect to account for individuals that were tracked for more than one year. We then combined both tracking periods and used the same model to determine population-level shifts in breeding ground arrival, nonbreeding ground departure, and migration duration across all years. To assess sex-specific differences at each stage, we re-ran each model with an interaction term between sex and year included among the predictor variables.

Next, we wanted to examine whether individuals tracked for multiple years exhibited trends that were reflective of population-level trends in breeding ground arrival and nonbreeding ground departure timing. Following Conklin et al. ^30^, we used subject-centering ^31^ to determine if there were differences in within- and between-individual trends. To do this, we used linear models with two variables – one for within individual variation, to ask if individuals with repeated measures were significantly shifting their migratory timing during their lifetime, and the other for between-individual variation, to ask if, over time, cohorts of individuals were shifting their migratory timing. These variables were then included as additive terms and arrival/departure date was used as the dependent variable. For these models, we only included tracks from multi-year individuals (n=19, tracks=46).

#### (b) Historic model of godwit arrival date

We had previously developed a statistical model that related weather variables from across the godwit migratory corridor with godwit first arrival dates in Anchorage, Alaska from 1974-2010 ^11^. That model identified six factors as influencing godwit migratory timing: (1) the previous 5-year mean May temperature from the breeding grounds, (2) the previous year’s first godwit arrival date at the breeding grounds, (3) the previous year’s mean May temperature from the breeding grounds, (4) the mean April wind speed from a northbound stopover site (Houston, TX), (5) the amount of precipitation during the nonbreeding season (Puerto Montt, Chile), and (6) the March value of the Southern Oscillation Index. This model had an R^2^ = 0.74. Here, we gathered data from the period since that paper was published (2011-2023) from the same data sources (www.ebird.org, www.ncdc.gov, and www.data.longpaddock.qld.gov.au) in order to predict the timing of godwit arrival in Beluga based on these environmental factors.

Using the *predict* function in ‘lme4’ with the previous historical model results ^11^ and the weather and eBird data from 2011-2023, we predicted first arrival dates for godwits over the same period for which we had geolocator data. We then ran a simple linear regression, with the predicted first arrival date as the dependent variable and year as the predictor variable, to derive an expected rate of change in godwit arrival dates since 2011.

## RESULTS

### Population-level shifts in migratory timing

There was no significant difference in either nonbreeding ground departure (*β* = -0.03, SE=0.49, p=0.96) or breeding ground arrival timing (*β* = -0.36, SE=0.57, p=0.53) for individuals tracked within the first tracking period (2010-2012). For individuals tracked within the second tracking period (2021-2023), there was no significant difference in nonbreeding ground departure timing (*β* = -0.95, SE=0.51, p=0.08), but breeding ground arrival advanced by ∼2.5 days over the 3-year period (*β* = -0.85, SE=0.30, p=0.01).

When the two tracking periods were combined, we found significant, directional, population-level shifts in both nonbreeding ground departure and breeding ground arrival timing across the 13 year-period (n=70). Godwits shifted both their nonbreeding ground departure (*β* = 0.36, SE =0.07; p<0.01; Fig. 1C) and breeding ground arrival timing (*β* = 0.33, SE=0.05; p<0.01; Fig. 1A) later by ∼6 days over the last 13 years. Despite observed changes in departure and arrival timing, migration duration stayed the same over this period (*β* = -0.02, SE=0.06; p=0.72; Fig. 1B), with individuals completing their northward migration in 24 days, on average.

**Figure 1.**
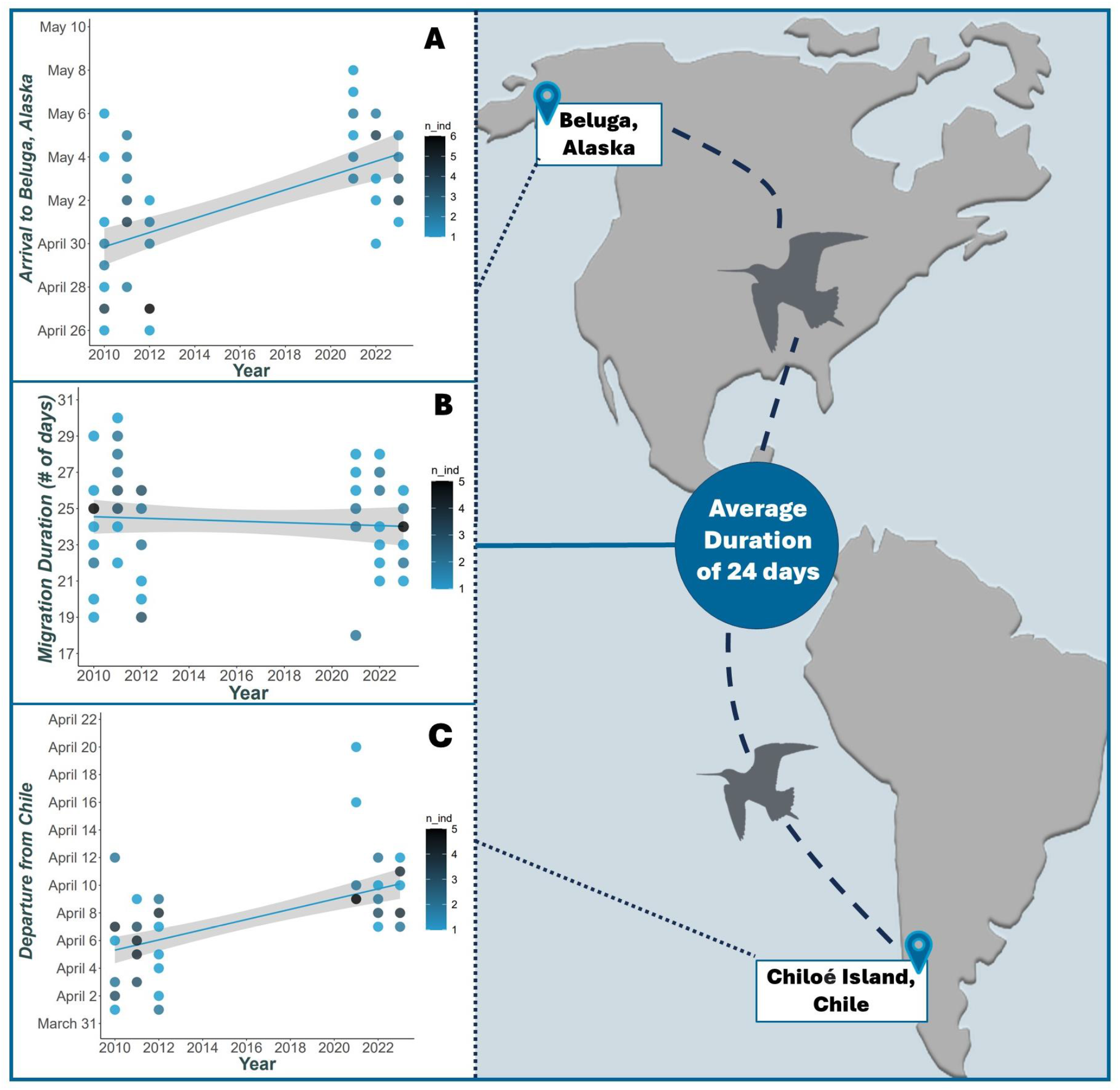
Changes in migratory timing for Hudsonian godwits breeding in Beluga, Alaska, USA. (A) Change in godwit population-level arrival timing to the breeding grounds in Beluga over a 13-year period (2010-2023); (B) Change in godwit population-level migration duration; (C) Change in godwit population-level departure timing from nonbreeding grounds in southern Chile. Points are scaled by color to represent the number of individuals with the same migration duration, departure, and arrival dates, respectively.

### Sex-specific shifts in migratory timing

We found no significant interaction between sex and year in population-level nonbreeding ground departure (interaction term: *β* = -1.96, SE=1.76; p=0.27), breeding ground arrival (interaction term: *β* = - 2.58, SE=1.41; p=0.07), or migration duration (interaction term: *β* = -0.02, SE=0.06; p=0.72).

### Shifts at the individual-level

For individuals that were tracked for multiple years, we found no significant difference in within-individual trends for either nonbreeding ground departure (*β* =0.69, SE=1.21, p=0.57; Fig. 2A & E) or breeding ground arrival timing (*β* =2.15, SE=1.14, p=0.07; Fig. 2B & F). However, between-individual trends were significantly different – both nonbreeding ground departure (*β* =1.63, SE=0.36, p<0.01) and breeding ground arrival (*β* =1.08, SE=0.34, p<0.01) shifted later between individuals. Only one individual, a female godwit (hereafter, “1PJ”), was tracked during both tracking periods – it shifted both its nonbreeding ground departure and breeding ground arrival timing later over time (Fig. 2C & D), aligning with our observed population-level trends (Fig. 1A & C).

**Figure 2.**
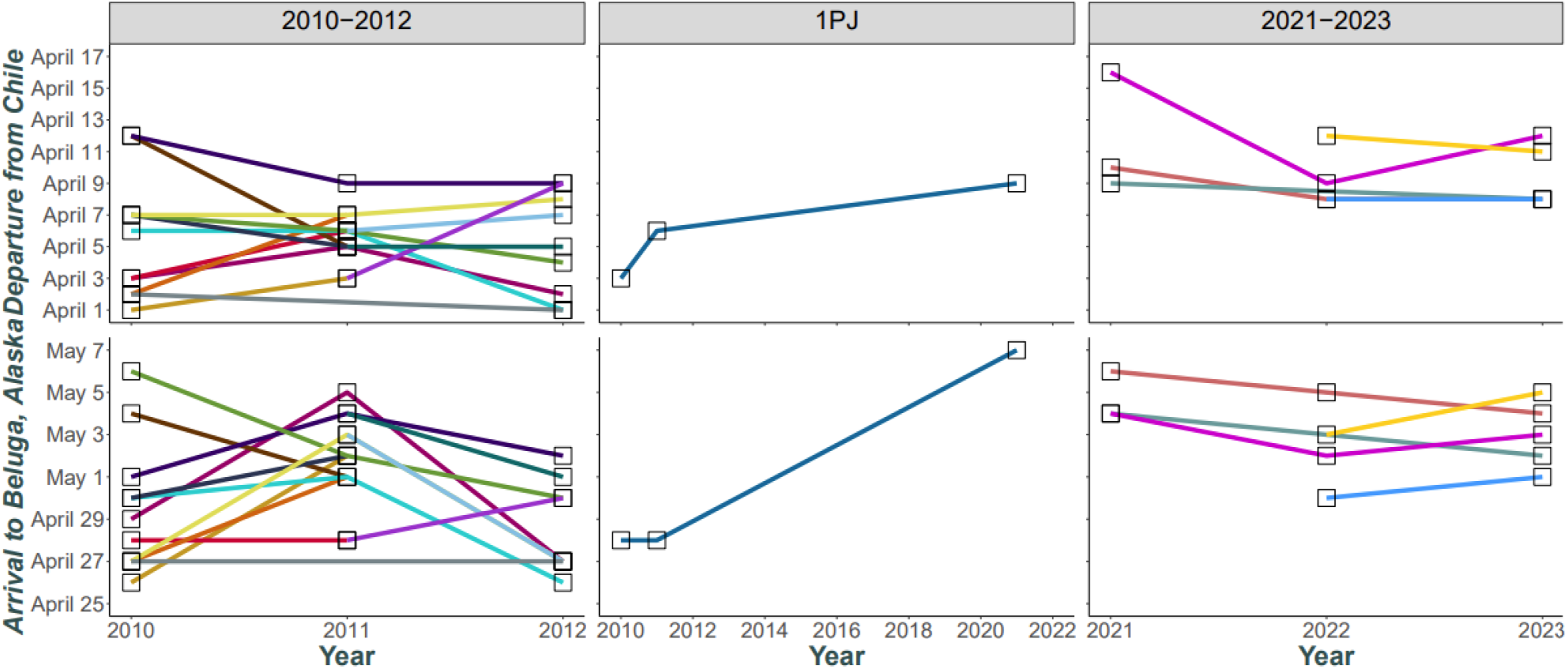
Variation in individual-level trends in migratory timing for Hudsonian godwits breeding in Beluga, Alaska, USA. (A) Nonbreeding ground departure and (B) breeding ground arrival timing from individuals tracked for either two or three years during the first tracking period, 2010-2012. (C) Nonbreeding ground departure and (D) breeding ground arrival timing for individual “1PJ”, which was tracked during both tracking periods. (E) Nonbreeding ground departure and (F) breeding ground arrival timing from individuals tracked for either two or three years during the second tracking period, 2021-2023. Colors represent unique individuals.

### Predictive model

Our historical model predicted no consistent change in population-level breeding ground arrival dates between 2011-2023 (*β* = -0.28, SE 0.15, p = 0.08), although there was a nonsignificant trend towards earlier breeding ground arrival (Fig. 3). However, actual first arrival dates derived from our geolocator data got significantly later during this period (*β* = 0.44, SE 0.11, p = 0.02; Fig. 3).

**Figure 3.**
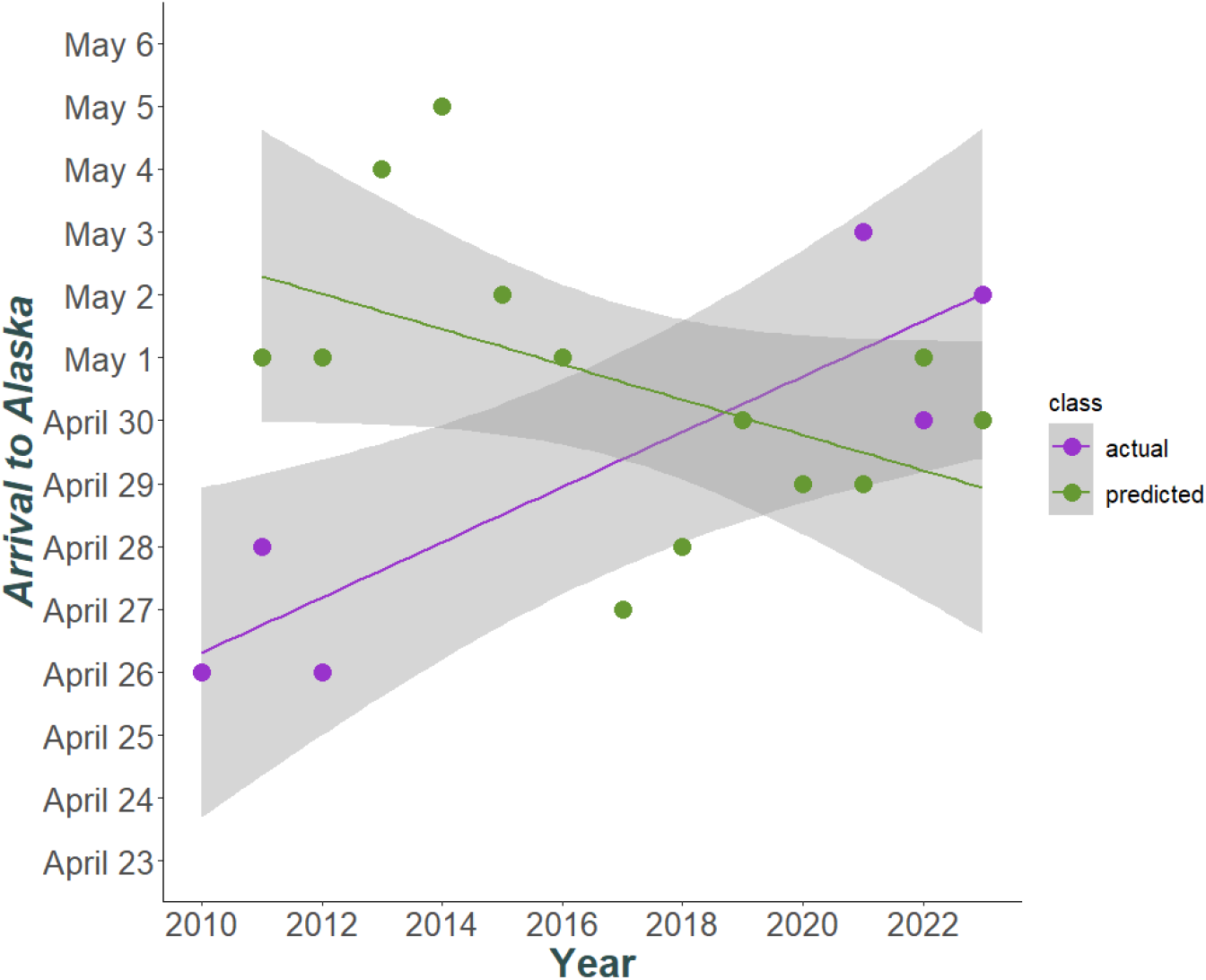
Trends in first arrival dates to Anchorage, Alaska, USA predicted with a historical model and actual, first arrival dates to Beluga, Alaska, USA derived from geolocators from 2010-2023.

## DISCUSSION

Contrary to the findings of many large-scale studies of long-distance migratory birds ^3,4,32^, we show that Hudsonian godwits, an extreme long-distance migratory shorebird can (1) rapidly alter their migratory timing, (2) use nonbreeding ground departure timing to drive changes in breeding ground arrival timing, and (3) dramatically reverse the direction in which their migratory timing is shifting within only a few years. Our results thus point to the need to better link individual- and species-level studies to understand both the patterns and drivers of migratory timing in the face of global change.

### Rapid change in population-level migratory timing

Previously, changes in godwit migratory timing of similar magnitude to the ones we observed here had occurred over a period of ∼40 years ^11^. Likewise, most large-scale studies that have assessed adjustments in migratory timing have found relatively slow rates of change over time – usually only 1-3 days per decade ^4,33,34^. However, we show that godwits have shifted their migratory timing by ∼6 days over a 13-year period – a remarkably fast rate of change. Previous large-scale studies have also frequently noted that short-distance migrants outpace long-distance migrants in the rate at which they have shifted their migratory timing ^33^. Yet, our results demonstrate that a species with one of the most extreme long-distance migrations of any bird species can alter its migratory timing at rates just as fast, if not faster, than short-distance migrants.

This rapid shift in godwit migratory timing suggests that individuals are capable of flexibly responding to environmental changes within their lifetime, something which has rarely, and only recently, been documented in other long-distance migrants ^30^. That said, in this study, we can only conclude that between-individual (i.e., intergenerational), and not within-individual (i.e., individual plasticity) trends are changing and contributing to population-level shifts in migratory timing over time. Many studies, like ours, do not have enough repeated individual measures over a sufficiently long period to differentiate the mechanistic role of between- and within-individual trends ^35,36^. As a result, deciphering whether developmental plasticity and evolutionary processes – or both simultaneously – are driving trends in migration timing remains a need in studies of responses to climate change ^35^. For example, individual plasticity has been found to account for only a fraction of responses to changing climatic trends, suggesting microevolution is taking place at the same time ^37^. Nevertheless, 1PJ, the one individual that we successfully tracked across our entire study period, attests to the possibility that godwits can exhibit dramatic flexibility in response to changes in climatic cues during their lifetimes (see also^30^). Therefore, with longer-term monitoring of repeatedly tracked individuals, we might expect to observe that both within- and between-individual trends can contribute to timing shifts at the population-level.

### Migratory timing shift driven by changes in departure timing

Historically, long-distance migrants have been predicted to be incapable of responding to environmental changes because of inherent constraints associated with their long-distance trips. For instance, the absence of cues predictive of breeding ground conditions on the nonbreeding grounds is thought to force individuals to predominantly rely on photoperiodic cues when initiating migration, making shifts in nonbreeding ground departure seem unlikely, if not, impossible ^7,8^. Further, the repeatability of nonbreeding ground departure timing in a plethora of avian species is relatively high ^38^. For these reasons, we predicted that godwits would achieve shifts in migratory timing primarily by utilizing *en route* strategies and that the two sexes would exploit strategies differently because of sex-specific goals.

Despite this, we found that godwits did not utilize *en route* strategies, as we observed no change in migration duration over time and no difference in migratory timing between the sexes. These findings support our prediction that females are limited in their ability to effectively utilize *en route* strategies because of the condition-dependent pressures of egg production ^17^. However, they contradict our prediction that males would utilize *en route* strategies, risking optimal body condition on arrival to achieve optimal arrival timing. This suggests that selection pressures may not be sex-specific but, rather, acting uniformly across the population. Nevertheless, sub-arctic and arctic breeding migrants are tasked with migrating vast distances in a matter of weeks and, in general, all individuals depart and arrive within a matter of days ^23^. Thus, if differences are present between the sexes, they may be small and difficult to distinguish with (relatively) small sample sizes.

Instead, contrary to past predictions ^7,8^ we observed a proportional shift, whereby nonbreeding ground departure changed in directional synchrony with arrival on the breeding grounds. To our knowledge, shifts of this magnitude have not been previously documented. For instance, bar-tailed godwits (*L. lapponica*) have been documented flexibly advancing their nonbreeding ground departure timing, but nonetheless failing to exhibit earlier arrival to the breeding grounds ^30^.

One possible explanation for this pattern in Hudsonian godwits is that individuals could be using knowledge of the previous year’s breeding ground conditions to inform their departure timing in subsequent years. While this concept is not novel ^9,11^, the predictive power of knowledge from previous years is likely low in highly stochastic climate regimes and only beneficial when breeding ground conditions are either unchanging or changing slowly, and linearly, over time. For instance, when godwits experienced slow, linear changes in climate in the past, information from previous breeding ground conditions strongly predicted godwit breeding ground arrival timing ^11^. The Arctic, however, is currently experiencing some of the most rapidly changing climate regimes ^39,40^ – causing godwits to experience increasingly variable conditions upon arrival in recent years ^22^. Thus, knowledge of previous years breeding ground conditions may be increasingly unreliable for godwits, limiting their ability to predict conditions within a given year and inform their subsequent migratory timing accordingly.

### Are godwits changing their migration timing adaptively?

Most studies have concluded that shifts in migratory timing occur in response to changes in climate ^3^. This largely precludes the possibility that shifts can be nonadaptive in nature or in response to other environmental conditions ^41^. Previously, godwits were advancing their breeding ground arrival timing and appeared to be shifting in relative synchrony with the climate ^11,21^. Using a model from these studies, we predicted that godwit migration timing should not have exhibited significant changes in recent years but, if anything, trend towards earlier breeding ground arrivals. However, in stark contrast, we found that over the last 13 years, godwits have significantly shifted their migratory timing 6 days later, suggesting that this may not be an adaptive response. We therefore propose an alternate hypothesis – new or different climatic and environmental factors on the nonbreeding grounds are imposing constraints on godwit migratory timing.

In southern Chile, the burgeoning aquaculture industry has changed many coastal areas, making coastal habitats increasingly dynamic. For instance, godwits, as well as other shorebirds, exclusively forage in the supratidal mudflats located in coastal bays that are also used for the cultivation and harvesting of fish, shellfish, and algae ^42–44^. The effects of such practices on shorebird communities are complex. In some cases, they may be mutually beneficial – areas with algae cultivation promote higher levels of benthic macroinvertebrates and are preferred by tactile-foraging shorebird species like godwits ^45^. In other cases, though, the human disturbance associated with the maintenance and harvesting of algae negatively influences godwit foraging efficiency and leads to decreases in body condition and survival in individuals from highly disturbed sites ^46–48^.

For godwits, the energetic expenditures and time associated with avoiding heavily disturbed areas could be especially costly in the days leading up to departure, which is immediately followed by a 7-day nonstop trans-Pacific flight ^23,49^. To sustain a flight of this magnitude, individuals must acquire substantial nutritional reserves in the weeks prior to departure ^50^. Godwits have thus been documented foraging throughout the day and night during this period ^51^. For this reason, disturbances may limit any surplus preparation time that godwits might possess ^47^. As a consequence, we might expect to see significant shifts, or in this case, delays, in migratory timing at the population-level.

Abiotic conditions, such as wind, could also influence migratory timing ^52^. For example, both advancements and delays in the timing of nonbreeding ground departure have been tied to the availability of beneficial tailwinds ^53^. Godwits that depart from southern Chile, in particular, are known to utilize prevailing tailwinds during their trans-Pacific flights ^49^. Thus, capitalizing on beneficial winds, or avoiding disadvantageous winds, is likely an important determinant of godwit migratory timing. It is possible that these winds are changing – favorable wind conditions may be occurring later in the season or becoming increasingly unpredictable due to changes in the Pacific subtropical anticyclone ^54,55^.

Individual godwits may therefore be energetically ready to depart, but remain grounded, delaying their departure until wind conditions are favorable. Because only wind conditions at a northbound stopover site were included in our historical model of breeding ground first arrival dates, we did not account for these potential trends in our predictions.

### Into the future

Taken together, we have documented an unexpected and rapid shift in migratory timing for an extreme long-distance migrant. While our evidence suggests that this shift may be driven by individual flexibility, we cannot make firm conclusions with our current sample size of repeated individual tracks. Nonetheless, differentiating between developmental plasticity, individual flexibility, and evolutionary processes is essential to assessing the rapidity with which migrants can respond to changes in their environments. Therefore, studies that employ a bottom-up approach – focusing on repeated individual-level measures – should be a high priority and coupled with larger scale, multi-species studies.

Our data also suggests that godwits are experiencing a constraint on their migratory timing on the nonbreeding grounds. We propose two potential factors that could be responsible for this pattern – increased anthropogenic disturbances and changing climatic conditions. Regardless, we emphasize that both factors have the potential to constrain entire populations from responding to climatic changes. Thus, both factors warrant future investigation. Further, delays in departure timing make stopover site quality another potential limiting factor, as degraded sites may prolong refueling duration^30^, in turn, further delaying breeding ground arrival, with the potential to subsequently affect reproductive success. Future studies should therefore focus on assessing and monitoring sites not just at the beginning and end of migratory journeys, but also in the middle. Currently, much of our knowledge regarding the vulnerability of long-distance migrants to climate change is derived from studies that only explore the influence of climatic factors on migratory timing and not other anthropogenic and environmental factors (but see ^56^). Looking to the future, we see the need for a more holistic approach – one that quantifies a diversity of variables from across the entire annual cycle – when assessing shifts in migratory timing and investigating their potential adaptiveness. For long-distance migrants, this holistic approach requires multi-hemispheric efforts and collaborations that, whilst ambitious, are imperative for addressing conservation concerns in a rapidly changing world.

## ACKNOWLEDGEMENTS

Thank you to Cook Inlet Region, Inc. for access to their traditional lands. Thank you also to the many field assistants that have helped in Beluga over the years. Finally, thank you to the Senner Lab for comments on earlier versions of this manuscript. Funding was provided by NSF-PCE 2318983 and 1110444, NSF DGE-1144153, USFWS-NMBCA 7320, the Association of Field Ornithologists, Wilson Ornithological Society, American Ornithological Society, Arctic Audubon Society, Cornell Lab of Ornithology, Athena Fund at the Cornell Lab of Ornithology, Faucett Family Foundation, David and Lucille Packard Foundation, and the University of South Carolina. Animal Care and Use approval was provided by the University of South Carolina (#2449-101417-042219) and University of Massachusetts Amherst (4136). Scientific approval was provided by the Alaska Department of Fish & Game (19-133, 20-024, 21-025, 22-033, and 23-113) and the U.S. Geological Survey (24-191).

## AUTHOR CONTRIBUTIONS

LP and NRS conceived of the project; all authors contributed to the data collection; LP carried out the analyses and wrote the manuscript; all authors contributed edits.

